# UBL3 Interacts with Alpha-synuclein in Cells and the Interaction is Downregulated by the EGFR Pathway Inhibitor Osimertinib

**DOI:** 10.1101/2023.05.15.540732

**Authors:** Bin Chen, Md. Mahmudul Hasan, Hengsen Zhang, Qing Zhai, A.S.M. Waliullah, Yashuang Ping, Chi Zhang, Soho Oyama, Mst. Afsana Mimi, Yuna Tomochika, Yu Nagashima, Tomohiko Nakamura, Tomoaki Kahyo, Kenji Ogawa, Daita Kaneda, Minoru Yoshida, Mitsutoshi Setou

## Abstract

Ubiquitin-like 3 (UBL3) acts as a post-translational modification (PTM) factor and regulates protein sorting into small extracellular vesicles (sEVs). sEVs have been reported as vectors for the pathology propagation of neurodegenerative diseases, such as α-synucleinopathies. Alpha-synuclein (α-syn) has been widely studied for its involvement in α-synucleinopathies. However, it is still unknown whether UBL3 interacts with α-syn, and is influenced by drugs or compounds. In this study, we investigated the interaction between UBL3 and α-syn, and any ensuing possible functional and pathological implications. We found that UBL3 can interact with α-syn by the *Gaussia princeps* based split luciferase complementation assay in cells and immunoprecipitation, while cysteine residues at its C-terminal, which are considered important as PTM factors for UBL3, were not essential for the interaction. The interaction was upregulated by 1-methyl-4-phenylpyridinium exposure. In drug screen results, the interaction was significantly downregulated by the treatment of osimertinib. These results suggest that UBL3 interacts with α-syn in cells and be significantly downregulated by epidermal growth factor receptor (EGFR) pathway inhibitor osimertinib. Therefore, the UBL3 pathway may be a new therapeutic target for α-synucleinopathies in the future.

## 1. Introduction

Ubiquitin-like 3 (UBL3), a highly conserved ubiquitin-like protein in eukaryotes, is localized to the cell membrane by prenylation [1]. The UBL3 gene is widely expressed in human tissues, with the strongest expression in the testis, ovary, and brain tissues [2]. In our previous study, UBL3 was characterized as a post-translational modification (PTM) factor regulating protein sorting to small extracellular vesicles (sEVs) [3]. Another recent study reported that UBL3 is involved in adaptive immunity by regulating ubiquitination-mediated transport of major histocompatibility complex II and CD86 [4]. The downregulated expression of UBL3 has been reported to be associated with human diseases, such as cervical cancer [5], gastric cancer [6], esophageal cancer [7], and non-small cell lung cancer [8]. Moreover, UBL3 can interact with more than 22 diseases-related proteins, including neurodegenerative disease-related molecules [3]. The previous proteomic analyses, however, may be insufficient since the results were only from MDA-MB-231 cells. Therefore, further exploration of the interactions between UBL3 and other proteins may facilitate the exploration of the potential effects of UBL3 and diseases.

Alpha-synuclein (α-syn), a highly conserved neuronal protein, is highly enriched in presynaptic nerve terminals. The physiological function of α-syn remains largely unclear, several biochemical activities have been proposed: regulation of dopamine neuro-transmission, and synaptic function/plasticity [9]. α-syn knockout mice do not exhibit a distinct phenotype [10]. Misfolded α-syn aggregated in α-synucleinopathies, such as Parkinson’s disease (PD), dementia with Lewy bodies [11], and multiple system atrophy [12]. Various factors are involved in the transition of α-syn from a physiological state to pathological aggregation, including genetic mutation [13], protein-protein interactions [14], PTM [15], and oxidative stress [16]. The phosphorylation of α-syn at Serine-129 may be important for the formation of inclusions in PD and related α synucleinopathies [17]. The interaction between α-syn and its protein interactomes play key roles in the pathological accumulation of α-syn. For example, α-syn interacts with synphilin-1 promoting the inclusion formation of α-syn [14]. α-syn interacts with molecular chaperone proteins [18] and protein deglycase DJ-1 [19] preventing the formation of oligomerized α-syn. Exploring potential proteins that can interact with α-syn is an essential direction to elucidate the aggregation mechanism of α-syn, and find new therapeutic targets for α-synucleinopathies. The pathological α-syn can be packaged in sEVs for cell-to-cell transport [20]. Until now, it is unknown whether α-syn interacts with UBL3.

Protein-protein interactions play essential roles in most biological processes [21] and offer a broad range of therapeutic targets for the treatment of human diseases [22]. Many natural products and drugs have been reported to be excellent molecule candidates for stabling or inhibiting protein-protein interactions [23]. The interaction between α-syn and the target protein can be affected by different compounds or drugs. Such as, the interaction between prolyl oligopeptidase and α-syn was downregulated by KYP-2047, a potent prolyl oligopeptidase inhibitor, reducing the accumulation of α-syn inclusion [24]. And, the protein-protein interaction of α-syn was downregulated by 03A10, a small molecule from the fruits of *Vernicia fordii* (Euphorbiaceae), inhibiting α-syn aggregation [25]. However, it remains unknown whether the interaction between UBL3 and α-syn will be affected by clinical drugs.

We hypothesise that UBL3 may interact with α-syn in cells and mediate the loading of α-syn into sEVs. In this study, we aim to investigate whether UBL3 can interact with α-syn in cells, and explore whether the treatment of clinical drugs affects the interaction between UBL3 and α-syn.

## 2. Materials and Methods

### 2.1. Animals

Age is an important risk factor for Parkinson’s disease and other neurodegenerative diseases. People usually develop the disease around age 60 or older [26]. Therefore, we choose elder mice with an average age of 22 months to investigate the level of phosphorylated Serine-129 (p-S-129). All of mice in this study were on a C57BL/6J background. Ubl3^-/-^ mice were acquired from the previously established laboratory colony [3]. The C57BL/6J strain wild type (WT) mice (SLC Inc., Hamamatsu, Shizuoka, Japan) were used as a control in this study. Three mice are included in each group. All mice were fed and bred at 12 hours of light/dark cycles. The genotypes of mice were confirmed by polymerase chain reaction (PCR) according to our previous report [3].

### 2.2. Antibodies and Drugs

Anti-p-S-129 α-syn antibody (Abcam, ab59264, 1:1000 dilution), anti-α-syn antibody (BioLegend, 834304, 1:1000 dilution), Anti-UBL3 antibody (ABclonal, A4028, 1:1000 dilution), anti-MYC antibody (MBL, M1932, 1:1000 dilution), anti-Flag antibody (MERCK, F7425-.2MG, 1:1000 dilution), horseradish peroxidase (HRP)-conjugated anti-rabbit secondary antibody (Cell signaling, 7074, 1:5000 dilution), biotinylated anti-rabbit immunoglobulin G (Vector Laboratories, BA-1000, 1:1000 dilution). All clinically approved drugs, natural products, and other bioactive components, a total of 32, were ordered from the corresponding suppliers (Supplemental Table 1) and used for drug screening. All drugs were dissolved in dimethyl sulfoxide (DMSO) to 10 mM stock solution.

### 2.3. Immunohistochemistry

Immunohistochemistry was conducted according to a previously published protocol [27]. Briefly, sections were incubated in 3% hydrogen peroxide in 1x phosphate-buffered saline (PBS, 0.1 mol/L, pH 7.4) for 15 minutes after serial deparaffinization, washed with 1x PBS three times, and treated with a solution containing 1% bovine serum albumin in 1x PBS for 1 hour at room temperature. Then samples were incubated with primary antibody for 1 hour at room temperature. After washing three times with 1x PBS, sections were treated for 1 hour with secondary antibody washed three times in 1x PBS, and processed using the avidin-biotin complex (Vector Laboratories, Newark, NJ, USA) and prepared in 1xPBS for 30 minutes at room temperature. The reaction was visualized using DAB (FUJIFILM Wako Pure Chemical Corporation, Osaka, Japan). Finally, the sections were subsequently counterstained using hematoxylin, dehydrated in graded alcohols (80%, 90%, 100%), transparentized with xylene, and coverslipped with PathoMount (FUJIFILM Wako Pure Chemical Corporation, Osaka, Japan). Images of immunohistochemistry were acquired using a NanoZoomer system (Hamamatsu Photonics, Hamamatsu, Shizuoka, Japan). Quantification of the immunoreactivity of representative areas was determined using the analysis software ImageJ (National Institutes of Health, Bethesda, MD, USA).

### 2.4. Plasmids Construction

Following the previous paper [28], for the the N-terminal region of *Gaussia princeps* luciferase (Gluc) sequence-tagged UBL3 (NGluc-UBL3) plasmid, the coding sequence of UBL3 (NM_007106) was inserted in the frame after the NGluc sequence in the pCI vector between the XhoI site and the MluI site. For the C-terminal region of Gluc sequence-tagged α-syn (α-syn-CGluc), the coding sequence of α-syn (NM_001146055.2) was inserted in the frame before the CGluc sequence in the pCI vector between the Xho I site and the Mlu I site. For the 3xFlag-UBL3 plasmid, the coding sequence of UBL3 was inserted in the frame after the 3xFlag sequence in the pcDNA3.1-3xFlag vector between the BamHI site and the EcoRI site. For the 6xMYC-α-syn plasmid, we amplified it by in vitro PCR using the corresponding primers. After digestion by XhoI and XbaI, the fragment of α-syn was inserted in the pcDNA3-6xMYC vector after the 6xMYC sequence. For the constructions of “CCVIL” amino acids in C-terminal region-deleted mutant of UBL3 (UBL3Δ5) UBL3Δ5, including NGluc-UBL3Δ5 and 3xFlag-UBL3Δ5, we deleted the last five amino acids (5’-CCVIL-3’) of NGluc-UBL3 and 3xFlag-UBL3 by in vitro site-directed mutagenesis using the corresponding primers (Table 1). The fragments of NGluc-UBL3, NGluc-UBL3Δ5, and α-syn-CGluc were tagged with an immunoglobulin kappa secretory signal (IKSS) sequence after the start codon. Sequences of all constructed plasmids were verified by Sanger sequencing. All primers are listed in Table 1.

**Table 1.**
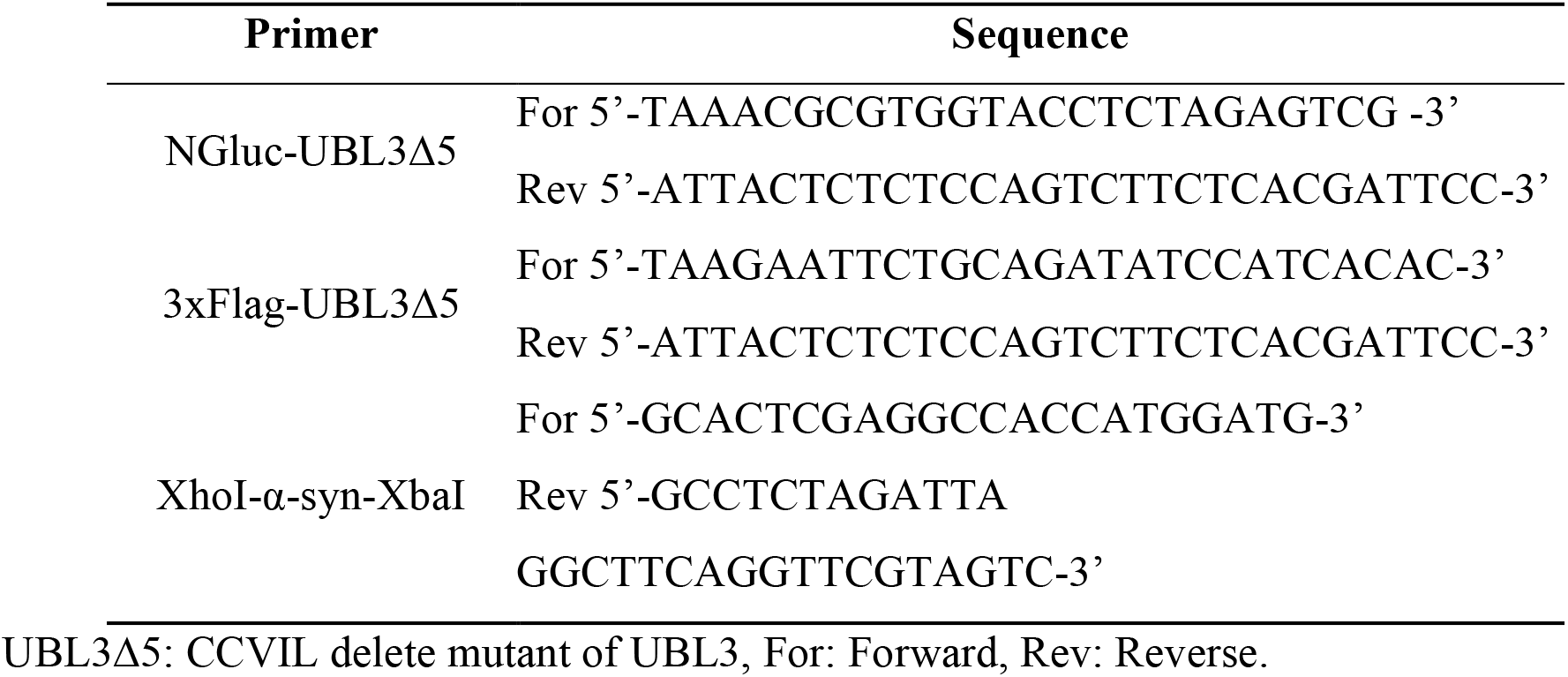
Primer list.

### 2.5. Cell culture and cDNA transfection

Human embryonic kidney (HEK) 293 cells (RIKEN Cell Bank, Tsukuba, Ibaraki, Japan) were cultured in Dulbecco’s modified Eagle’s medium (DMEM, Thermo Fisher Scientific, Waltham, MA, USA) with 10% fetal bovine serum (FBS). Cell cultures were incubated at 37 °C in a 5% CO_2_ humidified incubator. Cells were cultured in culture plates to 80%–90% confluence, and transiently transfected with cDNA plasmids using Lipofectamine 2000 transfect reagent (Thermo Fisher Scientific, Waltham, MA, USA) diluted in Opti-MEM reduced serum medium (Thermo Fisher Scientific, Waltham, MA, USA) according to the instruction of reagents.

### 2.6. Cell treatment

For the detection of UBL3 and α-syn interaction, the cell culture medium (CM) of transfected HEK293 cells was changed to FBS (-) Opti-MEM after transfected with NGluc-UBL3, NGluc-UBL3Δ5, and α-syn-CGluc plasmids for 12 hours. Then CM and cells were collected after further incubation for 3 days.

For the drug screening, HEK293 cells, after transfected using NGluc-UBL3 and α-syn-CGluc plasmids for 18 hours, were plated into 96-well cell culture plates in 100 μL/well of DMEM (10% FBS) and incubated overnight at 37 °C, in 5% CO_2_ humidified incubator. The next day, all solutions of candidate drugs were diluted in prewarmed DMEM (10% FBS) to 1.5 μM and 15 μM, 1.5-fold of the final concentration. Then 50 μL/well of the old culture medium was replaced with 100 μL of pre-warmed DMEM (10% FBS) containing the different candidate drugs to the final concentrations of 1 μM and 10 μM. For the treatment of MPP^+^, the solution of MPP^+^ (Cayman Chemical, Ann Arbor, MI, USA) was diluted in pre-warmed DMEM (10% FBS) to 1.5-fold of final concentrations and 50 μL/well of the old culture medium was replaced with 100 μL of pre-warmed DMEM (10% FBS) containing MPP^+^ to the final concentrations of 50 μM, 100 μM, 300 μM, 500 μM, and 600 μM, separately. The treatment of equivalent volume DMSO as the drug solution was set as a negative control. All cell culture plates were further incubated at 37 °C for another 48 hours. Then the CM was collected for luminescence intensity assay.

### 2.7. Sample preparation and Luciferase assay

All CM were centrifuged at 1200 rpm for 5 minutes to remove the cell debris. The collected HEK293 cells were lysed using cell lysis buffer (1% [v/v] Triton X100 in 1x PBS) and then centrifuged at 15000 rpm for 5 minutes to get supernatant. After adding 17 μg/ml coelenterazine (Cosmo Bio, Kyodo, Japan) diluted by Opti-MEM into CM and cell lysate (CL) supernatant, luminescence intensity was measured using a microplate reader (BioTek, Vermont, USA) immediately. The luminescence intensity of untreated DMEM (10% FBS) was set as a background. For the drug screening, all luminescence intensities of CM were corrected by subtracting the luminescence intensity of the background. The background corrected data was used to compute a ratio to the luminescence intensity compared to the background corrected luminescence intensity of DMSO.

### 2.8. 3-(4, 5-dimethylthiazol-2-yl)-2, 5-diphenyl-2H-tetrazolium Bromide (MTT) Assay

In live cells, MTT is a pale-yellow substrate that is cleaved by living cells to yield a dark blue formazan product. The amount of formazan reflects the number of living cells. For transfected HEK293 cells after collecting CM for the measurement of luminescence intensity, we added fresh pre-warmed DMEM (10% FBS) medium to 100 μL/well, mixed with 10 μL of MTT reagent (Sigma-Aldrich, Darmstadt, Hesse, Germany) to each well, and incubated in the incubator at 37 °C for 4 hours. After adding 100 μL/well of isopropanol with 0.04 N hydrogen chloride to dissolve the formazan, the absorbance (OD) of each well at a wavelength of 450 nm was detected using the microplate reader. Cell viability (OD _Intervention group_ – OD _Blank group_) / (OD _Control group_ – OD _Blank group_). The blank group had only medium without cells; the control group had medium and cells without intervention.

### 2.9. Bicinchoninic Acid (BCA) assay

According to the instructions of the BCA test kit (Thermo Fisher Scientific, Waltham, MA, USA), we add 20 μL of each standard or 2μL cell lysis sample diluted in 18μL water replicate into a microplate well. Add 200 μL of the working solution to each well and mix plate thoroughly on a plate shaker for 30 seconds. Incubate at 37°C for 30 minutes. Then measure the absorbance at 562 nm on a microplate reader after cool the plate to room temperature. Subtract the average absorbance measurement of the blank standard replicates from the measurements of all other individual standard and cell lysis sample replicates. Then calculate the protein concentration of each sample using the standard curve.

### 2.10. Co-Immunoprecipitation

For the validation of interaction between UBL3 and α-syn, HEK293 cells were transfected with 3xFlag-UBL3, 3xFlag-UBL3Δ5, and 6xMYC-α-syn plasmids in various combinations for 18 hours. After replacing with new CM, the transfected HEK293 cells were continually incubated for 36 hours, then washed and collected with ice-cold 1x PBS, pelleted by centrifugation at 1200 rpm for 5 minutes at 4 °C. Cell pellets were resuspended and lysed using 1% Triton lysate buffer (50 mM Tris-HCl [pH 7.4], 100 mM NaCl, and 1% [v/v] Triton X-100) for 30 minutes on ice. Cell debris and unbroken cells were removed by centrifugation at 15000 rpm for 15 minutes at 4 °C. Protein content of the supernatant was measured using the BCA assay according to the manufacturer’s instructions. Supernatant containing 500 μg of total protein were incubated with 50 μL of anti-Flag tag antibody magnetic beads (FUJIFILM Wako Pure Chemical Corporation, Osaka, Japan) with rotation for 10 hours at 4 °C. 50 μL of 2-mercaptoethanol (-) 2x sodium dodecyl sulfate (SDS) sample loading buffer (100 mM Tris-HCl [pH 6.8], 4% SDS, 20% glycerol, and 0.01% bromophenol blue) was added to the beads, after the beads were washed three times using ice-cold wash buffer (50 mM Tris-HCl [pH 7.4], 100 mM NaCl), and boiled at 95 °C for 5 minutes. CL and precipitated proteins were separated by SDS-PAGE for Western Blotting analysis.

### 2.11. Immunoblot

All samples, including CM and CL from transfected HEK293 cells and precipitated proteins, were loaded into 12% SDS-PAGE. Then the proteins were transferred to the polyvinylidene difluoride membrane. Membranes were blocked with shaking for 1 hour at room temperature using 0.5% [w/v] Skim milk in Tween-20 (+) Tris Buffered Saline (TBS-T; 100 mM Tris-HCl [pH 8.0], 150 mM NaCl, 0.5% [v/v] Tween-20), and then incubated with shaking overnight at 4 °C using the appropriate primary antibodies. The membranes were incubated with HRP-conjugated anti-rabbit secondary antibody with shaking at room temperature for 1 hour after three washes in TBS-T. The immunoreactive proteins were developed using the enhanced chemiluminescence kit (Thermo Fisher Scientific, Waltham, MA, USA) and detected on the FUSION FX imaging system (Vilber Lourmat, Collégien, Seine-et-Marne, France).

### 2.12. Statistical analysis

Measurement data were analyzed using GraphPad Prism 7.0 (GraphPad Software, LaJolla, CA, USA) statistical software, and expressed as mean ± SD (standard deviation). The differences between groups of data were calculated using Student’s t-test for unpaired data. P < 0.05 was considered statistically significant. All cell culture experiments were performed in triplicate.

## 3. Results

### 3.1. p-S-129 α-syn Is Upregulated in the Substantia Nigra of *Ubl3* Knockout (*Ubl3* ^-/-^) Mice

To explore whether UBL3 affects α-syn, the expression of p-S-129 α-syn was investigated in the brain tissues of the WT and *Ubl3*^-/-^ mice using anti-p-S-129 α-syn antibody by immunohistochemistry. The expression of p-S-129 α-syn was significantly upregulated (p = 0.0005) in the substantia nigra of *Ubl3*^-/-^ mice compared to WT mice (Figure 1), which suggests that UBL3 affects α-syn.

**Figure 1.**
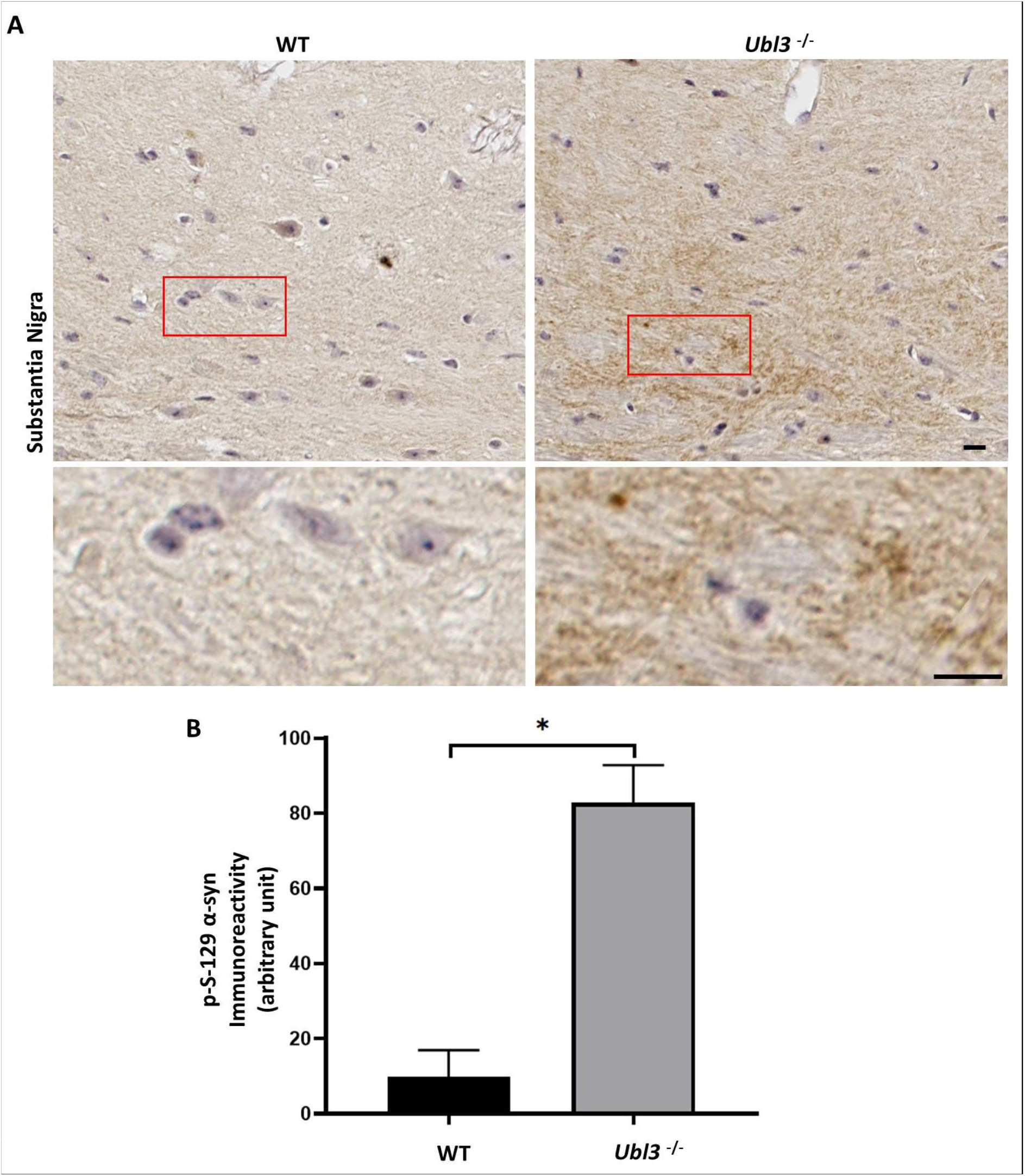
Expression of p-S-129 α-syn was upregulated in the substantia nigra of *Ubl3* ^-/-^ mice brain. (A) Representative images of immunocytochemistry staining of p-S-129 α-syn in the sub-stantia nigra of WT and *Ubl3*^-/-^ mice. Scale bars: 10 μm. (B) Quantification comparison of the im-munohistochemical expression of p-S-129 α-syn in the substantia nigra between WT and *Ubl3*^-/-^ mice. Histograms represent the mean + SD (the number of mice in each group was three). *: p-value < 0.05. WT: wild type. *Ubl3*^-/-^: *Ubl3* knock out.

### 3.2. UBL3 Interacts with α-syn in Cells

Further, we examined whether UBL3 interacts with α-syn using a *Gaussia princeps* based split luciferase complementation assay (SLCA), a powerful approach taking advantage of the reconstruction of the N-terminal and C-terminal fragments of Gluc to detect protein-protein interactions in vitro (Figure 2A) [29]. In this study, we constructed the NGluc-UBL3), NGluc-UBL3Δ5, and α-syn-CGluc plasmids. All constructions contain an IKSS sequence after the start codon (Figure 2B), which allows successfully expressed fragments and their interacting complexes in cells to be secreted into the CM.

**Figure 2.**
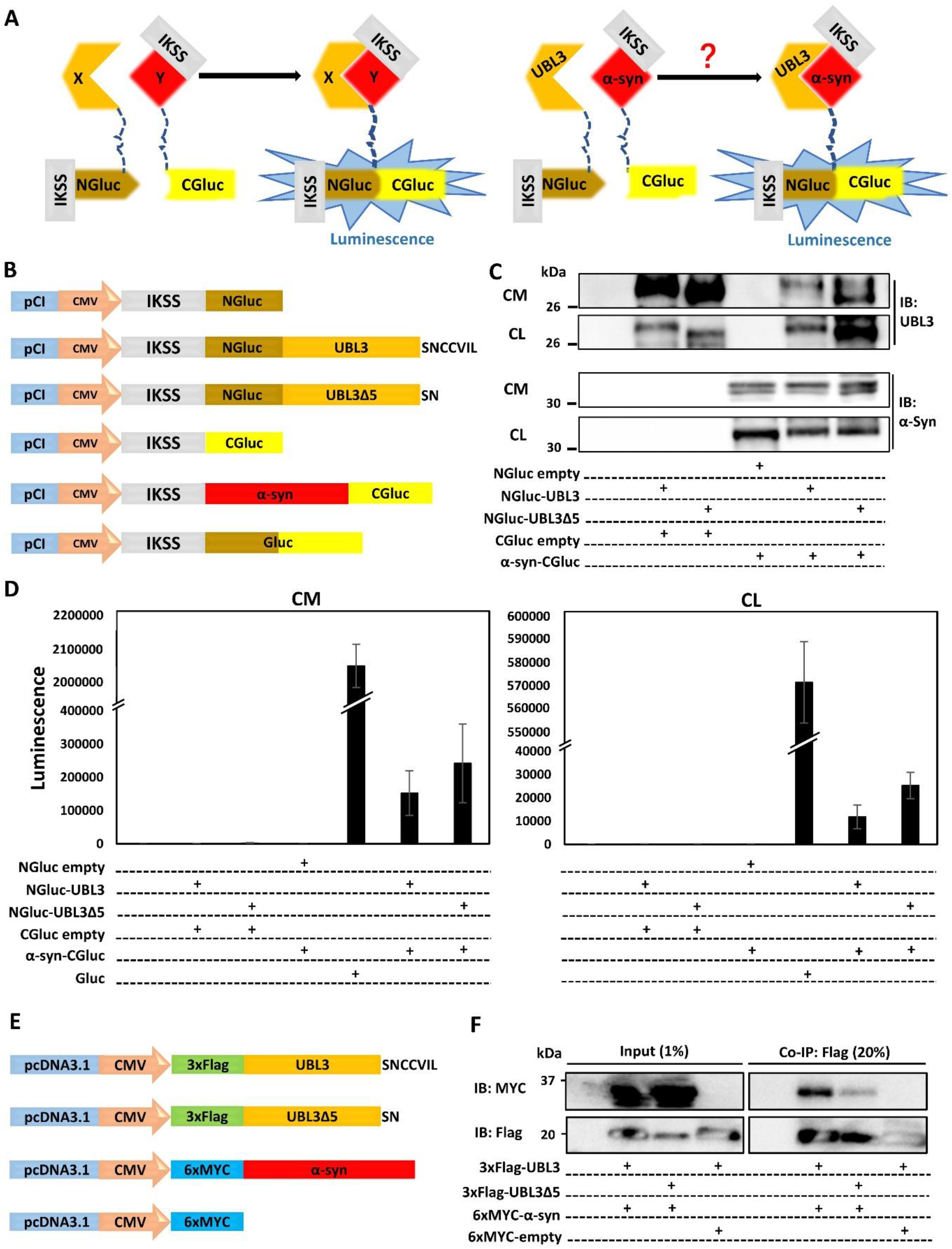
UBL3 interacts with α-syn in cells. (A) Schematic representation of the *Gaussia princeps* based split luciferase complementation assay system. (B) Schematic of split Gluc-tagged proteins, NGluc-UBL3, NGluc-UBL3Δ5, and α-syn-CGluc, both containing an IKSS sequence, which allows successfully expressed fragments and their interacting complexes to be secreted into the CM. (C) Immunoblotting of CM and CL from transfected HEK293 cells by anti-UBL3 polyclonal antibody and anti-α-syn antibody. (D) Luminescence of the CM (left) and CL (right) from transfected HEK293 cells. The luminescence ± SD in triplicate experiments is shown. (E) Schematic represen-tation of 3xFlag-UBL3, 3xFlag-UBL3Δ5, and 6xMYC-α-syn for co-immunoprecipitation (Co-IP). (F) Co-immunoprecipitated 3xFlag-UBL3 and 3xFlag-UBL3Δ5 interact with 6xMYC-α-syn. The input lanes are 1% of the sample prior to Co-IP, and the Co-IP lanes are 20% of the Co-IP products. IKSS: immunoglobulin kappa secretory signal; CM: cell culture medium; CL: cell lysate.

Then we co-expressed SLCA constructs in HEK293 cells in various combinations. Expressions of the SLCA constructs were confirmed by immunoblotting (Figure 2C). We measured the luminescence intensities of CM and CL from transfected HEK293 cells and found strong luminescence intensities from both fractions in the cells expressing NGluc-UBL3 + α-syn-CGluc, and NGluc-UBL3Δ5 + α-syn-CGluc. As a control, the CM and CL from Gluc overexpressing HEK293 cells showed markedly higher luminescence intensities compare to double-expression HEK293 cells. On the other hand, the luminescence intensities of CM and CL from HEK293 cells, expressing NGluc-UBL3, NGluc-UBL3Δ5, or α-syn-CGluc, were as low as the background level (Figure 2D).

To validate the interaction between UBL3 and α-syn, we constructed and co-expressed 3xFlag-UBL3, 3xFlag-UBL3Δ5 and 6xMYC-α-syn in various combinations in HEK293 cells (Figure 2E). The signal of 6xMYC-α-syn was detected from the co-immunoprecipitate of 3xFlag-UBL3 and 3xFlag-UBL3Δ5, while the signal of 6xMYC-α-syn in the coimmunoprecipitate of 3xFlag-UBL3Δ5 was less than that of 3xFlag-UBL3 (Figure 2F). These results showed that UBL3 interacts with α-syn in HEK293 cells.

### 3.3. The Interaction between UBL3 and α-syn in Cells Is Upregulated by the 1-methyl-4-phenylpyridinium (MPP^+^) Exposure

MPP^+^ has been reported to induce toxic aggregation of α-syn and the loss of nigrostriatal dopamine neurons [30,31]. To investigate whether the treatment of MPP^+^ affects the interaction between UBL3 and α-syn, we established an MPP^+^ treatment model using HEK293 cells transfected by NGluc-UBL3 and α-syn-CGluc cDNA. We collected CM and assayed the luminescence intensities after being treated with different concentrations of MPP^+^ for 48 hours, and also assessed the cell viability using an MTT assay. Although the luminescence intensities of CM from transfected HEK293 cells treated with different concentrations of MPP^+^ did not show significant differences (Figure 3A), the treatment with MPP^+^ at concentrations between 100 μM and 600 μM significantly inhibited cell viability in a concentration-dependent manner (Figure 3B). Therefore, to exclude the effect of cell activity on the luminescence intensities of CM, we divided the luminescence intensities by the cell viability to calculate the ratio of luminescence relative to the cell number. The treatment of MPP^+^ at concentrations between 300 μM and 600 μM significantly upregulated the interaction between UBL3 and α-syn in a dose-dependent manner (Figure 3C).

**Figure 3.**
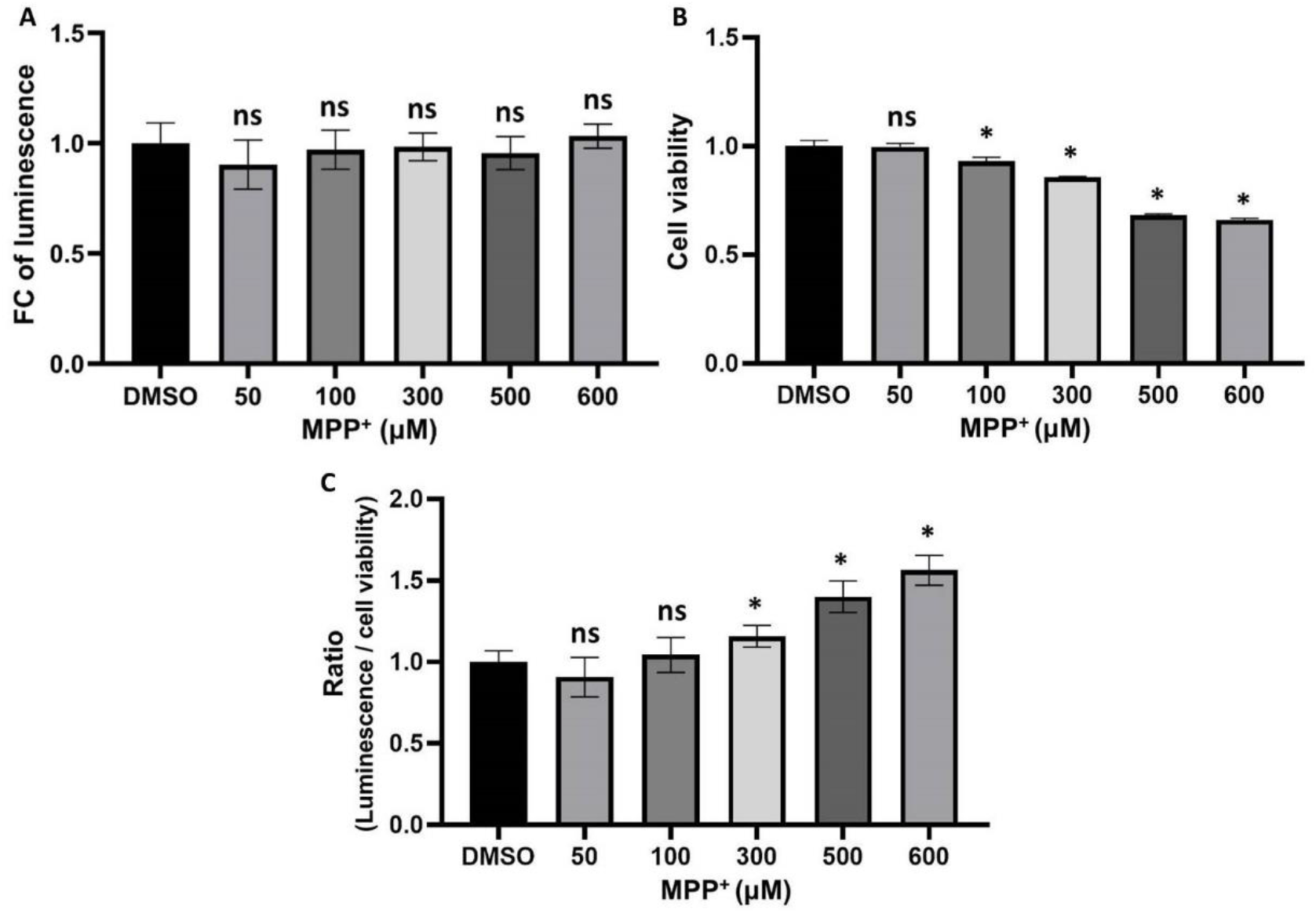
The interaction level in cells was upregulated by the treatment of MPP+. (A) Luminescence of CM from transfected HEK293 cells, which were treated with 50, 100, 300, 500, and 600 μM of MPP+ for 48 hours. (B) The cell viability of transfected HEK293 cells was treated with different concentrations of MPP+ for 48 hours. (C) The ratios of luminescence divided by cell viability were calculated in triplicate. The luminescence ± SD, cell viability ± SD, and ratio ± SD in triplicate are shown. ns (non-significant) p > 0.05, * p < 0.05. FC: Fold change. DMSO: dimethyl sulfoxide.

### 3.4. Interaction between UBL3 and α-syn in Cells Is Significantly Downregulated by Osimertinib

We used the HEK293 cells, transfected with NGluc-UBL3 and α-syn-CGluc cDNA, as a drug screening model to screen for drugs or compounds that can regulate the interaction between UBL3 and α-syn in cells. We assessed the luminescence intensities of CM from transfected HEK293 cells in the presence of 32 drugs (1 μM and 10 μM) at 48 hours in triplicate. All luminescence intensities of CM under drug treatment were normalized to that of vehicle treatment. Under the treatment with a concentration of 1 μM (Figure 4A), one drug, sulfasalazine, upregulated the luminescence intensities of CM by more than 25%. In contrast, one drug, docetaxel, downregulated the luminescence intensities of CM by more than 25%. Under the treatment with a concentration of 10 μM (Figure 4B), three drugs upregulated the luminescence intensities of CM by more than 25%, including sulfasalazine, pemetrexed, and gemcitabine. In contrast, four drugs downregulated the luminescence intensities of CM by more than 25%, including methylcobalamin, erlotinib, docetaxel, and osimertinib.

**Figure 4.**
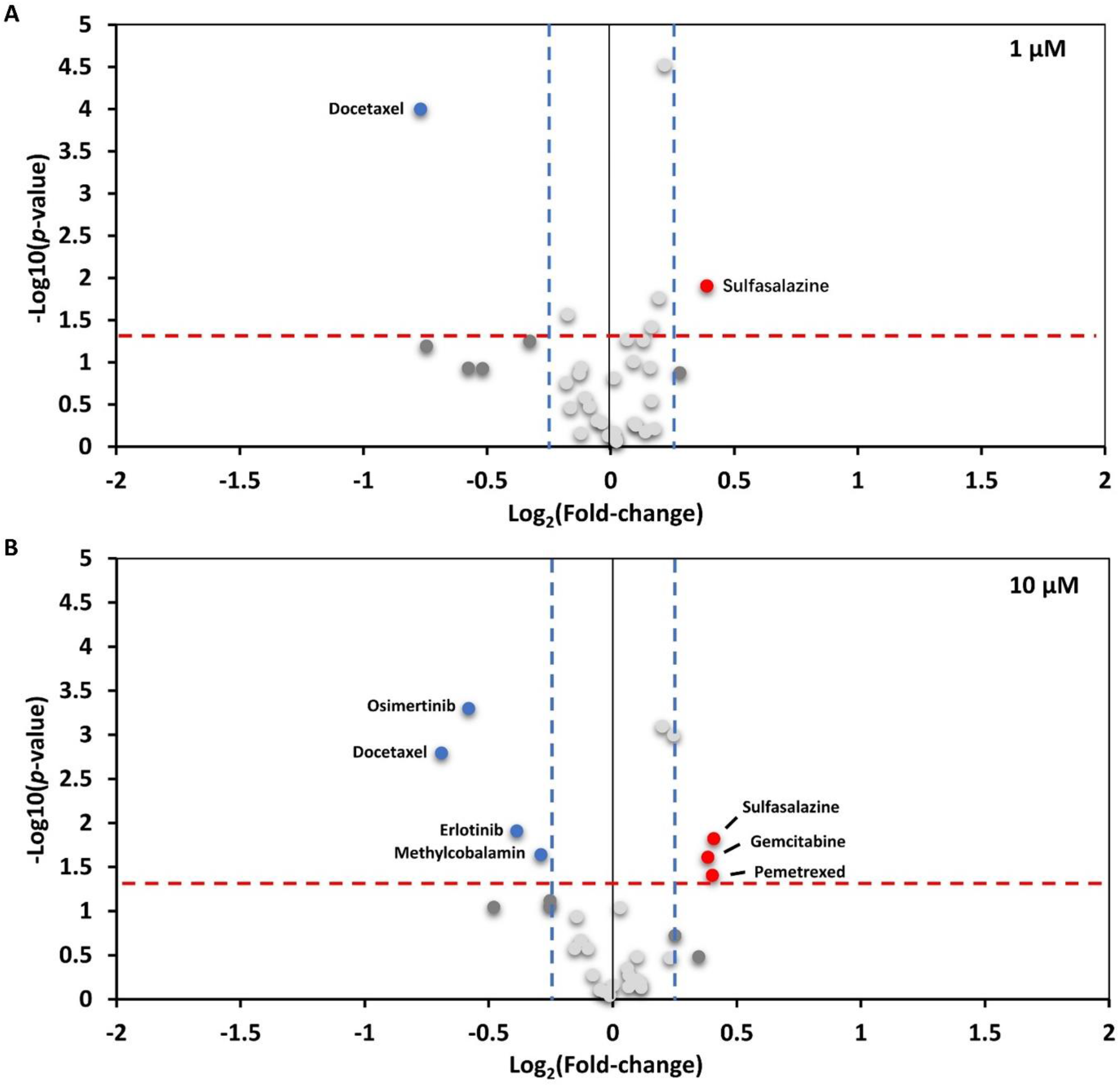
Drug screening results of 32 drugs at concentrations of 1 μM and 10 μM. (A) The screen result under a concentration of 1 μM. (B) The screen result under a concentration of 10 μM. The volcano plot showed the fold-change (x-axis) versus the significance (y-axis) of 32 drugs. The significance (non-adjusted *p*-value) and the fold-change are converted to −Log10(*p*-value) and Log2(fold-change), respectively. The vertical and horizontal dotted lines show the cut-off of fold-change ± 1.25, and *p*-value = 0.05, respectively. The luminescence of CM from transfected HEK293 cells was upregulated by > 1.25-fold with a *p*-value < 0.05 (upper-right, dots colored red), and the luminescence of CM from transfected HEK293 cells was downregulated by < -1.25-fold with *p*-value < 0.05 (upper-left, dots colored blue). The luminescence ± SD in triplicate experiments is shown. A student’s t-test was performed. *p* < 0.05 was considered statistically significant.

To exclude the effect of drug cytotoxicity on the drug screening results, we treated transfected HEK293 cells for 48 hours using selected drugs that significantly affected the interaction between UBL3 and α-syn at concentrations of 1 μM and 10 μM, and assessed the cell viability with an MTT assay. As shown in Figure 5A, the cell viability was significantly decreased by the treatment of docetaxel, gemcitabine, osimertinib, and pemetrexed at concentrations of 10 μM. Thus, the ratios of luminescence intensities of CM to cell viability were calculated (Figure 5B). At the concentration of 1 μM, the interaction between UBL3 and α-syn was upregulated by the treatment of gemcitabine (ratio = 1.37, p = 0.0008), while was significantly downregulated by the treatment of osimertinib (ratio = 0.83, p = 0.0021) and erlotinib (ratio = 0.73, p < 0.0001). At the concentration of 10 μM, the interaction was significantly upregulated the treatment of gemcitabine (ratio = 1.53 p = 0.015), while was significantly downregulated by the treatment of erlotinib (ratio = 0.72, p = 0.0002), and osimertinib (ratio = 0.55, p < 0.0001).

**Figure 5.**
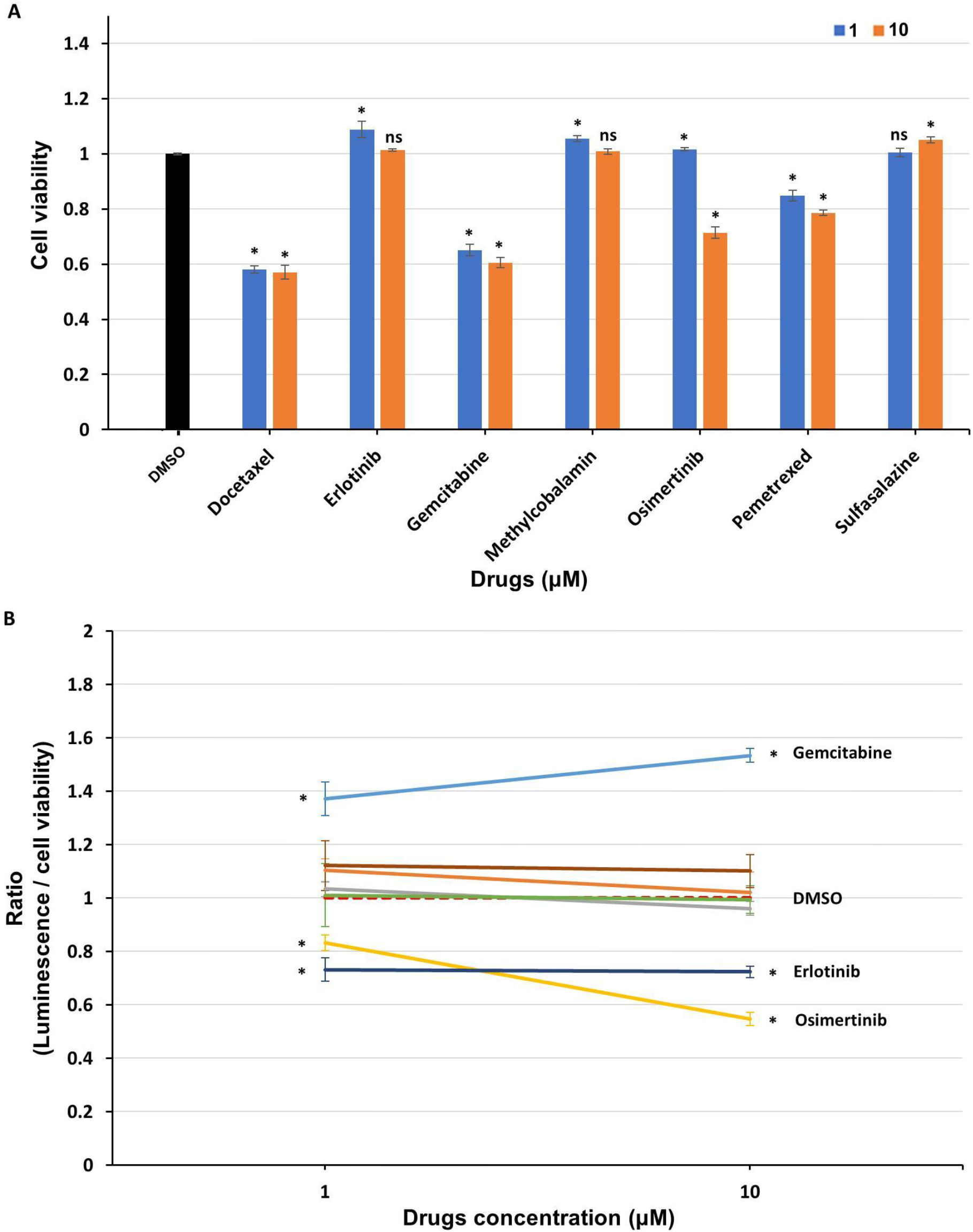
The interaction between UBL3 and α-syn in cells was regulated by the treatment of clinical drugs. (A) The cell viability of transfected HEK293 cells treated with selected drugs at concentrations of 1 and 10 μM for 48 hours, including docetaxel, erlotinib, gemcitabine, methylcobalamin, osimertinib, pemetrexed, and sulfasalazine. (B) The ratio of luminescence divided by cell viability was calculated in triplicate. The cell viability ± SD and the ratio ± SD in triplicate experiments are shown. A student’s t-test was performed. ns (non-significant) *p* > 0.05; * *p* < 0.05.

## 4. Discussion

This study first revealed that UBL3 could interact with α-syn in cells, and was upregulated in response to the MPP^+^ exposure. And the interaction could also be regulated by the treatment of clinical drugs. These results provided the first evidence that UBL3 may be involved in α-synucleinopathies with the possibility of being a potential therapeutic target.

Ageta et al. have reported that UBL3 can modify its protein interactomes through disulfide binding depending on the cysteine residues at its C-terminal [3]. It is interesting to note that our results showed that the interaction between UBL3 and α-syn in HEK293 cells is not completely erased after the deleting mutation of the cysteine residues at its C-terminus. UBL3, as a member of the ubiquitin-like protein family, contains a ubiquitin-like domain. Ubiquitin and ubiquitin-like proteins, such as NEDD8, SUMO, FAT10, and ISG15, are covalently attached to lysine residues of their protein interactomes through the C-terminal glycine residues [32]. Taken together, it is possible that, UBL3 interacts with α-syn in cells in another manner, rather than only dependent on cysteine residues at its C-terminal. In future studies, it is important to discover the interaction mechanism between UBL3 and α-syn.

Our results showed that the interaction between UBL3 and α-syn in cells was upregulated by the MPP^+^ exposure. MPP^+^, a bio-active derivative of 1-methyl-4-phenyl-1, 2, 3, 6-tetrahydropyridine, has been widely used as a common neurotoxin for both in vivo and in vitro experiments [33]. MPP^+^ has been reported as a key environmental risk factor of PD [30]. MPP^+^ exposure is known to disturb mitochondrial respiration by inhibiting the mitochondrial complex I, and this process plays a role in initiating mitochondrial dysfunction [34], which can induce and promote α-syn accumulation [35]. PD-like symptoms and aggregation of α-syn were observed in the MPP^+^-exposed rodent models [31]. Moreover, MPP^+^ exposure can also upregulate the expression of α-syn in SH-SY5Y cells [36]. Thus, the upregulation of interaction between UBL3 and α-syn induced by MPP^+^ exposure might be a response to the mitochondrial dysfunction and affect the accumulation of α-syn. This speculation will need to be investigated in the future.

The treatment of osimertinib downregulated the interaction between UBL3 and α-syn in cells significantly. Osimertinib, a third-generation epidermal growth factor receptor (EGFR) tyrosine kinase inhibitor, is widely used to treat non-small cell lung cancer [37]. In recent years, it has been suggested that the EGFR signaling pathway and associated genes possibly play an essential role in dopamine neuron cell death [38]. Exogenous neurotrophic supply of EGFR ligands rescues dopaminergic neurons from cell death induced by neurotoxins, 6-hydroxydopamine or MPTP [39]. An epidemiological study about the polymorphisms of the human EGFR gene found that rs730437 and rs11506105 polymorphisms of EGFR are possible in association with the susceptibility to PD [40]. The treatment of EGFR inhibitors can significantly reduce the p-S-129 α-syn pathology in mouse brain sections by reducing the level of seeding and propagation of pathological α-syn [41]. In our drug screening result, another RGFR inhibitor, erlotinib, also showed significant downregulation of interaction between UBL3 and α-syn. It is convincing from the view of the propagation pathway of α-syn pathology. α-syn can be secreted and transferred cell-to-cell via sEVs [20]. sEVs-associated α-syn can facilitate the propagation of α-syn pathology through cell-to-cell transfer [32]. And UBL3 interact with its target proteins and regulate the sorting of them into sEVs [3]. Therefore, these results indicated that interaction between UBL3 and α-syn may be associated with the inhibition of the α-syn pathology propagation via sEVs by crosstalk with the EGFR pathway.

In addition, we found that the treatment of gemcitabine also significantly upregulated the interaction between UBL3 and α-syn in cells. Gemcitabine, a nucleoside analog, has been widely used as an anticancer drug to treat a variety of cancers [42]. The activated gemcitabine triphosphate complex, formed by linking two phosphates, inhibits DNA synthesis by inhibiting ribonucleotide reductase [43]. The treatment of gemcitabine can induce the initiation of mitochondrial dysfunction [44]. This result is consistent with MPP^+^ exposure, suggesting that the upregulation of interaction between UBL3 and α-syn might be a response to the mitochondrial dysfunction.

This study had some limitations. Firstly, the exact mechanisms by which drug treatment affects the interaction between NGluc-UBL3 and α-syn-CGluc, altering protein expression, degradation, or directly influencing the process of interactions was not investigated. Secondly, whether the aggregation status of α-syn in HEK293 cells overexpressing α-syn affects the interaction between UBL3 and α-syn remains unstudied. And, due to technical constraints, it was not possible to determine whether the drug substance treatment would affect the activity of the luciferase itself. On the other hand, the impact of candidate drugs treatment and UBL3 itself to the aggregation state of α-syn is also not investigated. In the future, we will further explore these limitations in accordance with the methods summarized by previous report [45].

## 5. Conclusions

The results in this study showed that UBL3 interacts with α-syn in cells and the interaction between UBL3 and α-syn is upregulated in the response to MPP^+^ exposure. Moreover, it was downregulated by the treatment of EGFR inhibitor osimertinib. These findings provide the first evidence identifying UBL3 as an interacting protein of α-syn and the UBL3 may be a new therapeutic option for α-synucleinopathies in the future. This study extends the horizon for further etiological and therapeutic studies of α-synucleinopathies.

---

**Supplemental Table 1.** Drug list.

| **Drug** | **Supplier** | **Product ID** |
| --- | --- | --- |
| Apomorphine | Wako Bio | 013-18323 |
| Aniracetam | Combi-Blocks | OR-2762 |
| Chlorpromazine | TCI | C2481 |
| Chloroquine Diphosphate | Wako Bio | 038-17971 |
| Curcumin | Wako Bio | 038-04921 |
| Caffeic acid | Cayman Chemical Co. | 70602 |
| Cisplatin | Wako Bio | 033-20091 |
| Docetaxel | LKT Labs, Inc | D5709-10mg |
| Donepezil | Combi-Blocks | ST-7783 |
| Ergoloid | Cayman Chemical Co. | 24095 |
| Erlotinib | Med Chem Express | HY-50896 |
| Farnesol | Wako Bio | 064-03941 |
| Famotidine | Wako Bio | 066-06701 |
| Gefitinib | Wako Bio | 078-06561 |
| Gemcitabine | Combi-Blocks | QA-8591 |
| Haloperidol | Wako Bio | 084-04261 |
| Memantine hydrochloride | Combi-Blocks | OR-1164 |
| Melatonin | Wako Bio | 139-17111 |
| Mannitol | Wako Bio | 139-00842 |
| Methylcobalamin | Wako Bio | 138-14261 |
| Medetomidine Hydrochloride | Wako Bio | 139-17471 |
| Osimertinib | LC Laboratories | O-7200 |
| Pemetrexed | Combi-Blocks | QB-9701 |
| Pravastatin | Cayman Chemical Co. | 10010342 |
| Rivastigmine | Combi-Blocks | QC-5226 |
| Simvastatin | Fluorochem Ltd. | M03997 |
| Sulfasalazine | MP Biomedicals, Inc. | 191144 |
| Safinamide | Cayman Chemical Co. | S1472 |
| Scutellarin | Cayman Chemical Co. | 27461 |
| Trifluoperazine | TCI | T2849 |
| Tetrabenazine | Toronto Research Chemicals Inc | T284000 |
| Vitamin D2 | LKT Labs, Inc | V3476 |

## Reference

1. Downes, B.P.; Saracco, S.A.; Lee, S.S.; Crowell, D.N.; Vierstra, R.D. MUBs, a family of ubiquitin-fold proteins that are plasma membrane-anchored by prenylation. The Journal of biological chemistry 2006, 281, 27145–27157, doi:10.1074/jbc.M602283200.

2. Chadwick, B.P.; Kidd, T.; Sgouros, J.; Ish-Horowicz, D.; Frischauf, A.M. Cloning, mapping and expression of UBL3, a novel ubiquitin-like gene. Gene 1999, 233, 189–195, doi:10.1016/s0378-1119(99)00138-9.

3. Ageta, H.; Ageta-Ishihara, N.; Hitachi, K.; Karayel, O.; Onouchi, T.; Yamaguchi, H.; Kahyo, T.; Hatanaka, K.; Ikegami, K.; Yoshioka, Y.; et al. UBL3 modification influences protein sorting to small extracellular vesicles. Nature communications 2018, 9, 3936, doi:10.1038/s41467-018-06197-y.

4. Liu, H.; Wilson, K.R.; Firth, A.M.; Macri, C.; Schriek, P.; Blum, A.B.; Villar, J.; Wormald, S.; Shambrook, M.; Xu, B.; et al. Ubiquitin-like protein 3 (UBL3) is required for MARCH ubiquitination of major histocompatibility complex class II and CD86. Nature communications 2022, 13, 1934, doi:10.1038/s41467-022-29524-w.

5. Huang, L.; Zheng, M.; Zhou, Q.M.; Zhang, M.Y.; Yu, Y.H.; Yun, J.P.; Wang, H.Y. Identification of a 7-gene signature that predicts relapse and survival for early stage patients with cervical carcinoma. Medical oncology (Northwood, London, England) 2012, 29, 2911–2918, doi:10.1007/s12032-012-0166-3.

6. Shi, Y.; Qi, L.; Chen, H.; Zhang, J.; Guan, Q.; He, J.; Li, M.; Guo, Z.; Yan, H.; Li, P. Identification of Genes Universally Differentially Expressed in Gastric Cancer. BioMed research international 2021, 2021, 7326853, doi:10.1155/2021/7326853.

7. Singh, V.; Singh, L.C.; Vasudevan, M.; Chattopadhyay, I.; Borthakar, B.B.; Rai, A.K.; Phukan, R.K.; Sharma, J.; Mahanta, J.; Kataki, A.C.; et al. Esophageal Cancer Epigenomics and Integrome Analysis of Genome-Wide Methylation and Expression in High Risk Northeast Indian Population. Omics : a journal of integrative biology 2015, 19, 688–699, doi:10.1089/omi.2015.0121.

8. Zhao, X.; Yongchun, Z.; Qian, H.; Sanhui, G.; Jie, L.; Hong, Y.; Yanfei, Z.; Guizhen, W.; Yunchao, H.; Guangbiao, Z. Identification of a potential tumor suppressor gene, UBL3, in non-small cell lung cancer. Cancer biology & medicine 2020, 17, 76–87, doi:10.20892/j.issn.2095-3941.2019.0279.

9. 9. Winkler, S.; Hagenah, J.; Lincoln, S.; Heckman, M.; Haugarvoll, K.; Lohmann-Hedrich, K.; Kostic, V.; Farrer, M.; Klein, C. alpha-Synuclein and Parkinson disease susceptibility. Neurology 2007, 69, 1745–1750, doi:10.1212/01.wnl.0000275524.15125.f4.

10. Chandra, S.; Fornai, F.; Kwon, H.B.; Yazdani, U.; Atasoy, D.; Liu, X.; Hammer, R.E.; Battaglia, G.; German, D.C.; Castillo, P.E.; et al. Double-knockout mice for alpha- and beta-synucleins: effect on synaptic functions. Proceedings of the National Academy of Sciences of the United States of America 2004, 101, 14966–14971, doi:10.1073/pnas.0406283101.

11. Spillantini, M.G.; Schmidt, M.L.; Lee, V.M.; Trojanowski, J.Q.; Jakes, R.; Goedert, M. Alpha-synuclein in Lewy bodies. Nature 1997, 388, 839–840, doi:10.1038/42166.

12. Mezey, E.; Dehejia, A.; Harta, G.; Papp, M.I.; Polymeropoulos, M.H.; Brownstein, M.J. Alpha synuclein in neurodegenerative disorders: murderer or accomplice? Nature medicine 1998, 4, 755–757, doi:10.1038/nm0798-755.

13. Dawson, T.M.; Dawson, V.L. Molecular pathways of neurodegeneration in Parkinson’s disease. Science (New York, N.Y.) 2003, 302, 819–822, doi:10.1126/science.1087753.

14. Xie, Y.Y.; Zhou, C.J.; Zhou, Z.R.; Hong, J.; Che, M.X.; Fu, Q.S.; Song, A.X.; Lin, D.H.; Hu, H.Y. Interaction with synphilin-1 promotes inclusion formation of alpha-synuclein: mechanistic insights and pathological implication. FASEB journal : official publication of the Federation of American Societies for Experimental Biology 2010, 24, 196–205, doi:10.1096/fj.09-133082.

15. Schmid, A.W.; Fauvet, B.; Moniatte, M.; Lashuel, H.A. Alpha-synuclein post-translational modifications as potential biomarkers for Parkinson disease and other synucleinopathies. Molecular & cellular proteomics : MCP 2013, 12, 3543–3558, doi:10.1074/mcp.R113.032730.

16. Scudamore, O.; Ciossek, T. Increased Oxidative Stress Exacerbates α-Synuclein Aggregation In Vivo. Journal of neuropathology and experimental neurology 2018, 77, 443–453, doi:10.1093/jnen/nly024.

17. Smith, W.W.; Margolis, R.L.; Li, X.; Troncoso, J.C.; Lee, M.K.; Dawson, V.L.; Dawson, T.M.; Iwatsubo, T.; Ross, C.A. Alpha-synuclein phosphorylation enhances eosinophilic cytoplasmic inclusion formation in SH-SY5Y cells. The Journal of neuroscience : the official journal of the Society for Neuroscience 2005, 25, 5544–5552, doi:10.1523/jneurosci.0482-05.2005.

18. Burmann, B.M.; Gerez, J.A.; Matečko-Burmann, I.; Campioni, S.; Kumari, P.; Ghosh, D.; Mazur, A.; Aspholm, E.E.; Šulskis, D.; Wawrzyniuk, M.; et al. Regulation of α-synuclein by chaperones in mammalian cells. Nature 2020, 577, 127–132, doi:10.1038/s41586-019-1808-9.

19. Shendelman, S.; Jonason, A.; Martinat, C.; Leete, T.; Abeliovich, A. DJ-1 is a redox-dependent molecular chaperone that inhibits alpha-synuclein aggregate formation. PLoS biology 2004, 2, e362, doi:10.1371/journal.pbio.0020362.

20. Danzer, K.M.; Kranich, L.R.; Ruf, W.P.; Cagsal-Getkin, O.; Winslow, A.R.; Zhu, L.; Vanderburg, C.R.; McLean, P.J. Exosomal cell-to-cell transmission of alpha synuclein oligomers. Molecular neurodegeneration 2012, 7, 42, doi:10.1186/1750-1326-7-42.

21. Guo, W.; Wisniewski, J.A.; Ji, H. Hot spot-based design of small-molecule inhibitors for protein-protein interactions. Bioorganic & medicinal chemistry letters 2014, 24, 2546–2554, doi:10.1016/j.bmcl.2014.03.095.

22. Arkin, M.R.; Tang, Y.; Wells, J.A. Small-molecule inhibitors of protein-protein interactions: progressing toward the reality. Chemistry & biology 2014, 21, 1102–1114, doi:10.1016/j.chembiol.2014.09.001.

23. Fischer, G.; Rossmann, M.; Hyvönen, M. Alternative modulation of protein-protein interactions by small molecules. Current opinion in biotechnology 2015, 35, 78–85, doi:10.1016/j.copbio.2015.04.006.

24. Savolainen, M.H.; Yan, X.; Myöhänen, T.T.; Huttunen, H.J. Prolyl oligopeptidase enhances α-synuclein dimerization via direct protein-protein interaction. The Journal of biological chemistry 2015, 290, 5117–5126, doi:10.1074/jbc.M114.592931.

25. Wang, Q.; Yao, S.; Yang, Z.X.; Zhou, C.; Zhang, Y.; Zhang, Y.; Zhang, L.; Li, J.T.; Xu, Z.J.; Zhu, W.L.; et al. Pharmacological characterization of the small molecule 03A10 as an inhibitor of α-synuclein aggregation for Parkinson’s disease treatment. Acta pharmacologica Sinica 2023, doi:10.1038/s41401-022-01039-6.

26. Rademacher, D.J. Potential for Therapeutic-Loaded Exosomes to Ameliorate the Pathogenic Effects of & α-Synuclein in Parkinson & rsquo;s Disease. Biomedicines 2023, 11, 1187.

27. Guo, M.; Wang, J.; Zhao, Y.; Feng, Y.; Han, S.; Dong, Q.; Cui, M.; Tieu, K. Microglial exosomes facilitate α-synuclein transmission in Parkinson’s disease. Brain : a journal of neurology 2020, 143, 1476–1497, doi:10.1093/brain/awaa090.

28. Hashimoto, T.; Adams, K.W.; Fan, Z.; McLean, P.J.; Hyman, B.T. Characterization of oligomer formation of amyloid-beta peptide using a split-luciferase complementation assay. The Journal of biological chemistry 2011, 286, 27081–27091, doi:10.1074/jbc.M111.257378.

29. Wille, T.; Blank, K.; Schmidt, C.; Vogt, V.; Gerlach, R.G. Gaussia princeps luciferase as a reporter for transcriptional activity, protein secretion, and protein-protein interactions in Salmonella enterica serovar typhimurium. Applied and environmental microbiology 2012, 78, 250–257, doi:10.1128/aem.06670-11.

30. Tieu, K. A guide to neurotoxic animal models of Parkinson’s disease. Cold Spring Harbor perspectives in medicine 2011, 1, a009316, doi:10.1101/cshperspect.a009316.

31. Sonsalla, P.K.; Zeevalk, G.D.; German, D.C. Chronic intraventricular administration of 1-methyl-4-phenylpyridinium as a progressive model of Parkinson’s disease. Parkinsonism & related disorders 2008, 14 Suppl 2, S116–118, doi:10.1016/j.parkreldis.2008.04.008.

32. Welchman, R.L.; Gordon, C.; Mayer, R.J. Ubiquitin and ubiquitin-like proteins as multifunctional signals. Nature reviews. Molecular cell biology 2005, 6, 599–609, doi:10.1038/nrm1700.

33. Lotharius, J.; O’Malley, K.L. The parkinsonism-inducing drug 1-methyl-4-phenylpyridinium triggers intracellular dopamine oxidation. A novel mechanism of toxicity. The Journal of biological chemistry 2000, 275, 38581–38588, doi:10.1074/jbc.M005385200.

34. Kim, H.Y.; Jeon, H.; Kim, H.; Koo, S.; Kim, S. Sophora flavescens Aiton Decreases MPP(+)-Induced Mitochondrial Dysfunction in SH-SY5Y Cells. Frontiers in aging neuroscience 2018, 10, 119, doi:10.3389/fnagi.2018.00119.

35. Grünewald, A.; Kumar, K.R.; Sue, C.M. New insights into the complex role of mitochondria in Parkinson’s disease. Progress in neurobiology 2019, 177, 73–93, doi:10.1016/j.pneurobio.2018.09.003.

36. Lu, M.; Sun, W.L.; Shen, J.; Wei, M.; Chen, B.; Qi, Y.J.; Xu, C.S. LncRNA-UCA1 promotes PD development by upregulating SNCA. European review for medical and pharmacological sciences 2018, 22, 7908–7915, doi:10.26355/eurrev_201811_16417.

37. Tan, C.S.; Gilligan, D.; Pacey, S. Treatment approaches for EGFR-inhibitor-resistant patients with non-small-cell lung cancer. The Lancet. Oncology 2015, 16, e447–e459, doi:10.1016/s1470-2045(15)00246-6.

38. Kim, I.S.; Koppula, S.; Park, S.Y.; Choi, D.K. Analysis of Epidermal Growth Factor Receptor Related Gene Expression Changes in a Cellular and Animal Model of Parkinson’s Disease. International journal of molecular sciences 2017, 18, doi:10.3390/ijms18020430.

39. Gill, S.S.; Patel, N.K.; Hotton, G.R.; O’Sullivan, K.; McCarter, R.; Bunnage, M.; Brooks, D.J.; Svendsen, C.N.; Heywood, P. Direct brain infusion of glial cell line-derived neurotrophic factor in Parkinson disease. Nature medicine 2003, 9, 589–595, doi:10.1038/nm850.

40. Jin, J.; Xue, L.; Bai, X.; Zhang, X.; Tian, Q.; Xie, A. Association between epidermal growth factor receptor gene polymorphisms and susceptibility to Parkinson’s disease. Neuroscience letters 2020, 736, 135273, doi:10.1016/j.neulet.2020.135273.

41. Tavassoly, O.; Del Cid Pellitero, E.; Larroquette, F.; Cai, E.; Thomas, R.A.; Soubannier, V.; Luo, W.; Durcan, T.M.; Fon, E.A. Pharmacological Inhibition of Brain EGFR Activation By a BBB-penetrating Inhibitor, AZD3759, Attenuates α-synuclein Pathology in a Mouse Model of α-Synuclein Propagation. Neurotherapeutics : the journal of the American Society for Experimental NeuroTherapeutics 2021, 18, 979–997, doi:10.1007/s13311-021-01017-6.

42. Pandit, B.; Royzen, M. Recent Development of Prodrugs of Gemcitabine. Genes 2022, 13, doi:10.3390/genes13030466.

43. Cerqueira, N.M.; Fernandes, P.A.; Ramos, M.J. Understanding ribonucleotide reductase inactivation by gemcitabine. Chemistry (Weinheim an der Bergstrasse, Germany) 2007, 13, 8507–8515, doi:10.1002/chem.200700260.

44. Inamura, A.; Muraoka-Hirayama, S.; Sakurai, K. Loss of Mitochondrial DNA by Gemcitabine Triggers Mitophagy and Cell Death. Biological & pharmaceutical bulletin 2019, 42, 1977–1987, doi:10.1248/bpb.b19-00312.

45. Estaun-Panzano, J.; Arotcarena, M.L.; Bezard, E. Monitoring α-synuclein aggregation. Neurobiology of disease 2023, 176, 105966, doi:10.1016/j.nbd.2022.105966.

